# Contribution of pretomanid to novel regimens containing bedaquiline with either linezolid or moxifloxacin and pyrazinamide in murine models of tuberculosis

**DOI:** 10.1101/514661

**Authors:** Jian Xu, Si-Yang Li, Deepak V. Almeida, Rokeya Tasneen, Kala Barnes-Boyle, Paul J. Converse, Anna M. Upton, Khisimuzi Mdluli, Nader Fotouhi, Eric L. Nuermberger

## Abstract

Novel regimens combining bedaquiline and pretomanid with either linezolid (BPaL regimen) or moxifloxacin and pyrazinamide (BPaMZ regimen) shorten the treatment duration needed to cure TB in BALB/c mice compared to the first-line regimen and have yielded promising results in initial clinical trials. However, the independent contribution of the investigational new drug pretomanid to the efficacy of BPaMZ has not been examined and its contribution to BPaL has been examined only over the first 2 months of treatment. In the present study, the addition of pretomanid to BL increased bactericidal activity, prevented emergence of bedaquiline resistance, and shortened the duration needed to prevent relapse with drug-susceptible isolates by at least 2 months in BALB/c mice. Addition of pretomanid to BMZ resulted in a 1 log_10_ greater CFU reduction after 1 month of treatment and/or reduced the number of mice relapsing in each of 2 experiments in BALB/c mice and in immunocompromised nude mice. Bedaquiline-resistant isolates were found at relapse in only one BMZ-treated nude mouse. Treatment of infection with a pyrazinamide-resistant mutant in BALB/c mice with BPaMZ prevented selection of bedaquiline-resistant mutants and reduced the proportion of mice relapsing compared to BMZ alone. Among severely ill C3HeB/FeJ mice with caseous pneumonia and cavitation, BPaMZ increased median survival (≥60 vs. 21 days) and reduced median lung CFU by 2.4 log_10_ at 1 month compared to BMZ. In conclusion, in 3 different mouse models, pretomanid contributed significantly to the efficacy of the BPaMZ and BPaL regimens, including restricting the selection of bedaquiline-resistant mutants.

---

The World Health Organization (WHO) estimates that 10.4 million people developed active tuberculosis (TB) in 2016 and 1.67 million people died from it (1). Nearly 500,000 new cases of multidrug-resistant (MDR) TB occur annually, with an estimated treatment success rate of only 54% (1, 2). The current standard short-course regimen for drug-susceptible TB consisting of rifampin (RIF), isoniazid (INH), pyrazinamide (PZA), and ethambutol (EMB) (regimen abbreviated as RHZE) requires a 6-month treatment duration to provide sufficient population-level efficacy. It takes 9-24 months for regimens containing at least 4-6 drugs, including at least one injectable agent, to treat patients with MDR-TB (3). New regimens to shorten and simplify TB treatment are urgently needed. If such regimens do not contain INH or RIF, they may be applicable to both drug-susceptible and MDR-TB.

The combination of bedaquiline (BDQ) + pretomanid (PMD) + moxifloxacin (MXF) + PZA (regimen abbreviated as BPaMZ) had superior bactericidal and sterilizing activity compared to RIF+INH+PZA in a murine model of TB, shortening the duration of treatment required to prevent relapse by 2.5-3.5 months (4). In the subsequent phase 2 NC-005 trial (NCT02193776), PZA-susceptible MDR-TB patients receiving the BPaMZ regimen had significantly faster sputum culture conversion than drug-susceptible TB patients receiving RIF+INH+PZA+EMB (5), suggesting that the results in mice may translate well to the clinic. A phase 3 trial evaluating the BPaMZ regimen administered for 4 months in drug-susceptible TB patients and for 6 months in MDR-TB patients is now enrolling subjects (NCT03338621). The combination of BDQ + PMD + linezolid (LZD) (regimen abbreviated as BPaL) also has superior bactericidal and sterilizing activity compared to RHZE in a murine TB model (6). Although it does not cure mice as rapidly as BPaMZ, this regimen has a greater spectrum of activity and has recently shown promising efficacy as an all-oral 6-month regimen in patients (Nix-TB trial) with extensively drug-resistant TB (7).

It is important to understand the contribution of each component in a regimen that is moving forward in the clinic. The independent contributions of BDQ, MXF and PZA to the efficacy of BPaMZ were previously demonstrated in a BALB/c mouse TB model (4). Furthermore, receipt of the BPaMZ regimen was associated with numerically higher sputum conversion rates in PZA-susceptible MDR-TB patients compared to drug-susceptible TB patients receiving the BDQ+PMD+PZA regimen and PZA-resistant MDR-TB patients receiving BPaMZ in the NC-005 trial (5), indicating the contribution of MXF and PZA, respectively. In addition, the sputum conversion rates after 2 months of treatment with BPaMZ in the NC-005 trial were higher than those in MDR-TB patients receiving the same regimen without BDQ in the NC-002 trial (8), indicating the contribution of BDQ.

PMD is a nitroimidazole drug that is activated within *Mycobacterium tuberculosis* by the bacterial deazaflavin-dependent nitroreductase Ddn and has bactericidal activity against replicating and non-replicating bacilli (9, 10). The contribution of this investigational new drug to the BPaMZ regimen has yet to be confirmed directly in pre-clinical or clinical studies. Indeed, addition of PMD antagonized the bactericidal activity of BDQ, BDQ+PZA and BDQ+PZA+clofazimine (CFZ) in past experiments in mice (11–13). However, the addition of PMD increased the bactericidal activity when added to BDQ+LZD and increased both the bactericidal and sterilizing activity when added to BDQ+sutezolid (6, 12). Another possible advantage of including PMD in the BPaMZ regimen is that it could reduce the selection of BDQ-resistant mutants, since such mutants would be more effectively targeted by PaMZ (PMD + MXF + PZA) than by MZ (MXF + PZA) alone (since PaMZ is a synergistic combination (14)), when considering the reliably active drugs remaining in the regimen.

The present study was undertaken to confirm the independent contributions of PMD to BPaMZ and BPaL by assessing the efficacy of each regimen with and without inclusion of PMD. Both regimens were evaluated in the same high-dose aerosol infection model in BALB/c mice in which their therapeutic potential was first described (4, 6). The contribution of PMD to the 4-drug BPaMZ regimen was further evaluated in athymic nude mouse and C3HeB/FeJ mouse models of TB. Athymic nude mice, which lack mature, differentiated T cells and are thus deprived of cell-mediated immunity, are more prone to relapse and the emergence of drug-resistant mutants than BALB/c mice (15, 16), thereby providing a more stringent model for evaluating a regimen’s ability to truly sterilize the infection and/or prevent the selection of drug-resistant mutants. This model may be more representative of TB in patients with immunocompromising diseases such as human immunodeficiency virus (HIV) or following iatrogenic immunosuppression who have an increased risk of treatment failure and relapse, especially those not receiving antiretroviral therapy (17).

C3HeB/FeJ mice are increasingly being utilized for TB drug development because, unlike BALB/c mice which develop only cellular granulomas following infection with *M. tuberculosis*, the former develop caseating necrotic lung lesions, including cavities, that more closely resemble the pathological hallmarks of human TB (18–21). The necrotic nature of these lesions can alter drug partitioning and present different microenvironments at the site of infection in various lesion sub-compartments that affect the overall efficacy of some drugs (18, 19, 22–24). Therefore, comparative studies are useful to evaluate the potential impact of these pathological differences on drug efficacy.

## Results

### Experiment 1. Comparison of BPaMZ and BMZ in BALB/c and athymic nude mice

The scheme of this experiment is shown in Table S1. The aerosol infection implanted nearly 4 log_10_ CFU in the lungs of both mouse strains (Table 1). At the start of the treatment 14 days post-infection, BALB/c mice and nude mice harbored approximately 7.95 and 7.56 log_10_ CFU in their lungs, respectively. After 1 month of treatment, the addition of PMD to BMZ resulted in an additional reduction of approximately 1 log_10_ in both BALB/c and nude mice (*P* < 0.01). Irrespective of the regimen, the decrease in lung CFU counts was significantly greater in BALB/c mice compared to nude mice, with additional 1.1 log_10_ reductions observed with both regimens (*P* < 0.01). Based on prior experience (4), BALB/c mice were expected to be culture-negative after 2 months of treatment and were not assessed at that time point. Among nude mice, all mice in the BPaMZ group and 7 of 10 mice in the BMZ group were culture-negative at 2 months despite plating the entire lung homogenate. Only a few CFU were detected in the other 3 mice of the BMZ group.

**Table 1.**
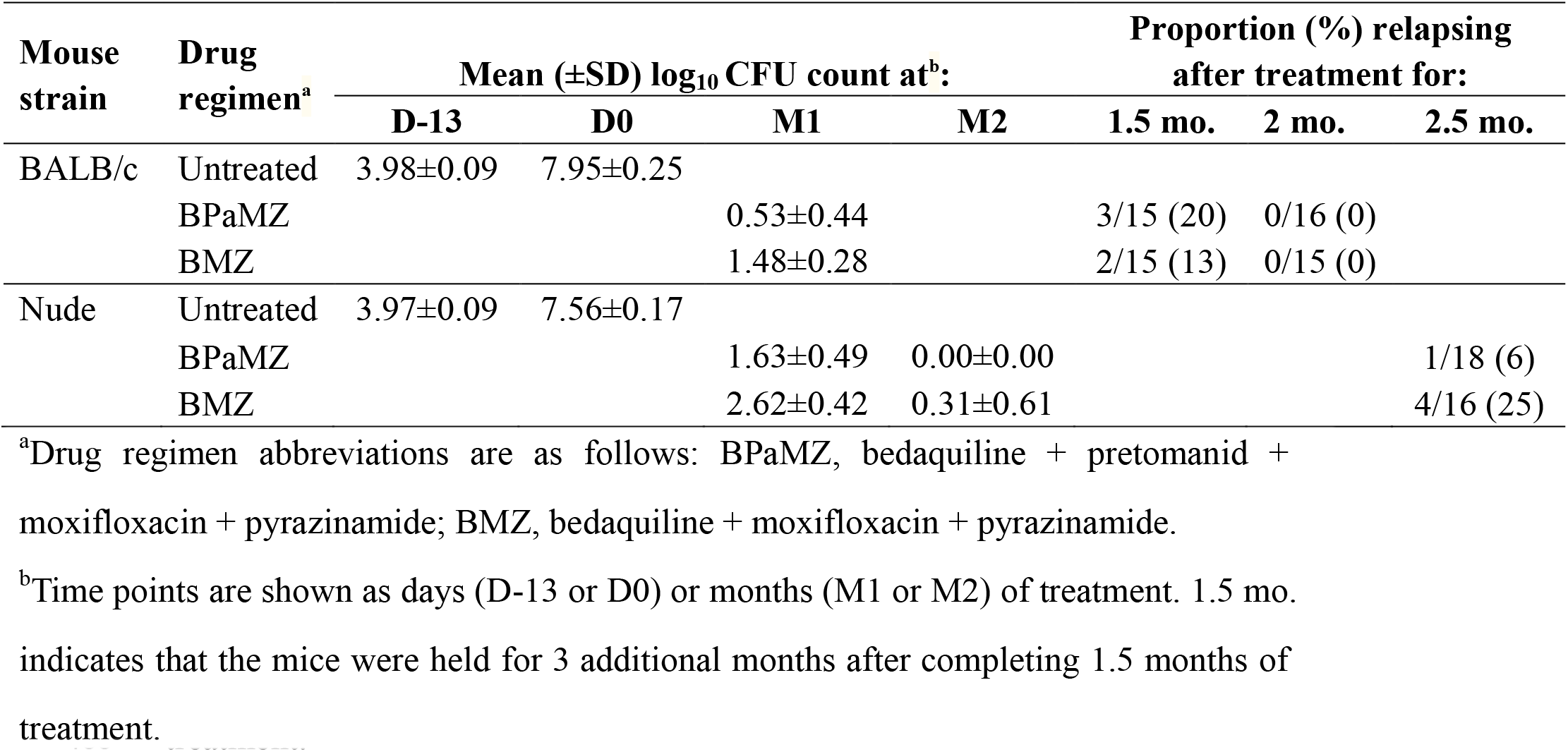
Lung CFU counts assessed during treatment and proportion of mice relapsing after treatment completion in Experiment 1

Relapse was assessed 3 months after completing 1.5 and 2 months of treatment in BALB/c mice and after 2.5 months of treatment in nude mice (Table 1). In BALB/c mice, no significant difference was observed in the proportions of mice relapsing after treatment with BPaMZ (3/15 [20%)] or BMZ (2/15 [13%]) for 1.5 months (*P* = 1.0). Both groups were relapse-free after 2 months of treatment. Among nude mice, which required a longer duration of treatment to cure, the proportion relapsing after 2.5 months of treatment was higher in BMZ-treated mice (4/16 [25%]) compared to BPaMZ-treated mice (1/18 [6%]), but the difference was not statistically significant (*P* = 0.16). Three nude mice in the BMZ group were euthanized when they became moribund just 6 weeks after completing treatment. Colonies isolated from lung homogenates were confirmed to be *M. tuberculosis* by colony morphology, acid-fast staining and 16s rRNA sequencing. Therefore, these mice were counted as relapses. Three nude mice in the BPaMZ group also required euthanasia when they became moribund 9 weeks after completing treatment. However, the lung homogenates from these 3 mice yielded no growth except 1 colony from the lungs of one mouse that was subsequently identified by colony morphotype, AFB staining, and 16s rRNA sequencing as *Staphylococcus epidermidis*. Therefore, these mice were not counted as relapses.

We hypothesized that the addition of PMD to BMZ would reduce the selection of BDQ-resistant mutants in nude mice. At the start of treatment, lung homogenates from 5 mice were plated in parallel on media containing 0.06 μg/ml of BDQ or 2 μg/ml of PMD. The mean frequencies of CFU isolated on BDQ- and PMD-containing plates were 1.3 × 10^-6^ and 6.1 × 10^-6^, respectively, among the total CFU counted on drug-free plates. Three to five individual BDQ-resistant colonies were selected from each mouse for sequencing of the *Rv0678, pepQ*, and *atpE* genes. Spontaneous *Rv0678* mutants were identified in all 5 mice and unique *pepQ* mutants were also found in 2 of the 5 mice (Table S2). None of the 7 colonies tested had *atpE* mutations. In total, 15 unique mutations (most of them frameshift mutations) were scattered across the *Rv0678* gene. Two mutants isolated on BDQ-containing plates (colonies 8 and 16) were selected for whole genome sequencing (WGS) to confirm the mutations. WGS confirmed the *Rv0678* mutations previously identified by PCR-based sequencing and the absence of other mutations. Among relapsing mice, a single colony grew on BDQ-containing plates from one of 3 nude mice relapsing 6 weeks after completing 10 weeks of BMZ treatment. It harbored a c313t (R105C) mutation in *rv0678*. However, this mutant represented a very small proportion of the total CFU count similar to the baseline frequency at D0, suggesting that it reflected a spontaneous mutation arising during multiplication after treatment ended. Growth amounting to more than 1% of the total CFU count was observed on BDQ-containing plates from one mouse relapsing at 12 weeks post-treatment with BMZ and sequencing revealed a *pepQ* mutation (g896t), indicating that this mutant was likely selectively amplified during treatment. Among all the BDQ-resistant mutants, frameshift mutations in *Rv0678* were routinely associated with 2-fold-higher MICs than single nucleotide polymorphisms (SNPs) in *Rv0678* and *pepQ* (Table 2).

**Table 2.** Lung CFU counts assessed during treatment against *M. tuberculosis* H37Rv (wild-type) and a *pncA* mutant (in italics) and proportion of mice relapsing after treatment completion in Experiment 2

| Regimen<sup>a</sup> | Mean lung log<sub>10</sub> CFU count (±SD)<sup>b</sup> |  |  |  | Proportion of mice relapsing after treatment for<sup>c</sup>: |  |  |  |  |
| --- | --- | --- | --- | --- | --- | --- | --- | --- | --- |
|  | D-14 | D0 | M1 | M2 | 1 mo. | 1.5 mo. | 2 mo. | 3 mo. | 4 mo. |
| Untreated | 4.06±0.05 | 7.90±0.16 |  |  |  |  |  |  |  |
| BL |  |  | 4.87±0.16 | 2.69±0.30 |  |  | 15/15<sup>7</sup> | 15/15<sup>5</sup> | 14/15<sup>5</sup> |
| BPaL |  |  | 3.29±0.09 | 0.68±0.24 |  |  | 7/15 | 0/15 | 0/15 |
| BMZ |  |  | 1.29±0.19 |  | 15/15 | 6/15 | 1/15 |  |  |
| BPaMZ |  |  | 1.05±0.18 |  | 14/15 | 0/15 | 0/15 |  |  |
| *Untreated* | 4.36±0.17 | 8.09±0.08 |  |  |  |  |  |  |  |
| *BMZ* |  |  | 4.06±0.23 | 1.24±0.17 |  |  | 15/15<sup>3</sup> | 7/20<sup>3</sup> |  |
| *BPaMZ* |  |  | 4.22±0.23 | 1.61±0.32 |  |  | 15/15<sup>1</sup> | 0/20 |  |

<sup>a</sup>Drug regimen abbreviations are as follows: BL, bedaquiline + linezolid; BPaL, bedaquiline + pretomanid + linezolid; BMZ, bedaquiline + moxifloxacin + pyrazinamide; BPaMZ, bedaquiline + pretomanid + moxifloxacin + pyrazinamide.
<sup>b</sup>Time points are shown as days (D-14 or D0) or months (M1 or M2) of treatment. 1 mo. indicates that the mice were held for 3 additional months after completing 1 month of treatment.
<sup>c</sup>Superscripts represent the number of mice with isolates resistant to 0.125 mg/L BDQ.

### Experiment 2. Confirmation of the contribution of PMD to the BPaMZ and BPaL regimens in BALB/c mice and evaluation of the impact of baseline PZA- or PMD-resistance

A second experiment was performed to confirm the results of Experiment 1, to evaluate the contribution of PMD to BPaMZ in the event of baseline PZA resistance, to assess the contribution of PMD to the sterilizing activity of BPaL, and to confirm that the contribution of PMD to BPaMZ and BPaL is directly attributable to its anti-tuberculosis activity upon activation by Ddn. The schemes for this experiment are shown in Tables S3 and S4. Mice were infected in parallel with the H37Rv strain or either of the isogenic PZA- or PMD-resistant mutants. Mean lung CFU counts exceeded 4 log_10_ CFU on the day after infection (Table 2). At the start of the treatment 2 weeks later, mean CFU counts were approximately 8 log_10_ CFU in the H37Rv and *pncA* mutant infection groups, respectively, and modestly lower in the *ddn* mutant group. Among mice infected with the H37Rv parent, the addition of PMD to BMZ did not result in a statistically significant decrease in lung CFU counts after 1 month of treatment. However, the addition of PMD was associated with lower CFU counts at the relapse assessments after 1 (*P* = 0.001) and 1.5 months of treatment (*P* = 0.02), as well as fewer relapses after 1.5 months of treatment (*P* = 0.02) (Figure 1, Table 2). The isolate from the mouse that relapsed after 2 months of BMZ treatment was not BDQ-resistant.

**Figure 1.**
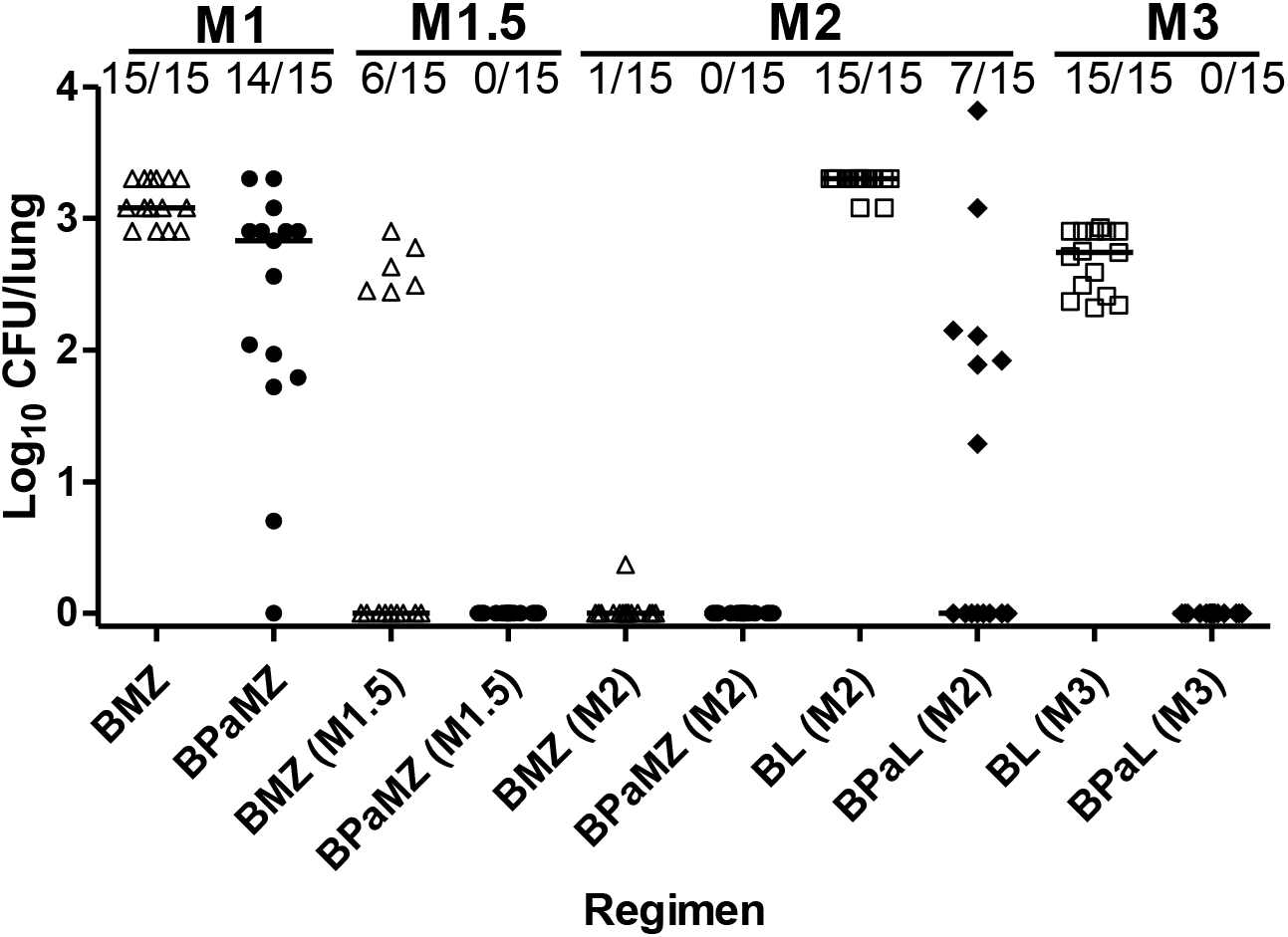
Proportion of relapses and individual mouse lung CFU counts after treatment of infection with *M. tuberculosis* H37Rv for one month (M1), one-and-a-half months (M1.5), two months (M2), and three months (M3) with each regimen. Regimen symbols: BMZ open triangles, BPaMZ solid circles, BL open squares, and BPaL solid diamonds. Horizontal black line indicates the median.

As expected, both regimens were significantly less effective against the *pncA* mutant, consistent with the important contribution of PZA previously observed in wild-type infections (4). No significant effect of PMD was observed in the mean CFU counts after 1 or 2 months of treatment (Table 2). However, addition of PMD was associated with significantly fewer relapses (*P* = 0.01) (Table 2) and lower CFU counts at the relapse assessment (*P* = 0.01) after 3 months of treatment (Figure 2). Addition of PMD also prevented the selection of BDQ-resistant mutants in *pncA* mutant-infected mice. Seven mice receiving BMZ (3 and 4 mice treated for 2 and 3 months, respectively) had growth on plates containing BDQ 0.125 μg/ml plates that exceeded 1% of the growth on drug-free plates (range, 15-100%), compared to just one mouse receiving BPaMZ for 2 months (*P* = 0.05). Six of the 7 isolates tested from BDQ-containing plates had mutations in *Rv0678* (5) or *pepQ* (1) (Table 3). After 2 months of treatment in *pncA* mutant-infected mice, 3/15 relapses in the BMZ group had BDQ-resistant CFU with unique mutations of g362a and an a436 insertion in *Rv0678* and a g812 insertion in *pepQ* vs. 1/15 in the BPaMZ group with an a202g mutation in *Rv0678*. After 3 months of treatment, 3 relapses in the BMZ group were BDQ-resistant CFU with t407c substitution or a g168 deletion in *Rv0678* in 2 isolates and wild-type *Rv0678* and *pepQ* sequences.

**Figure 2.**
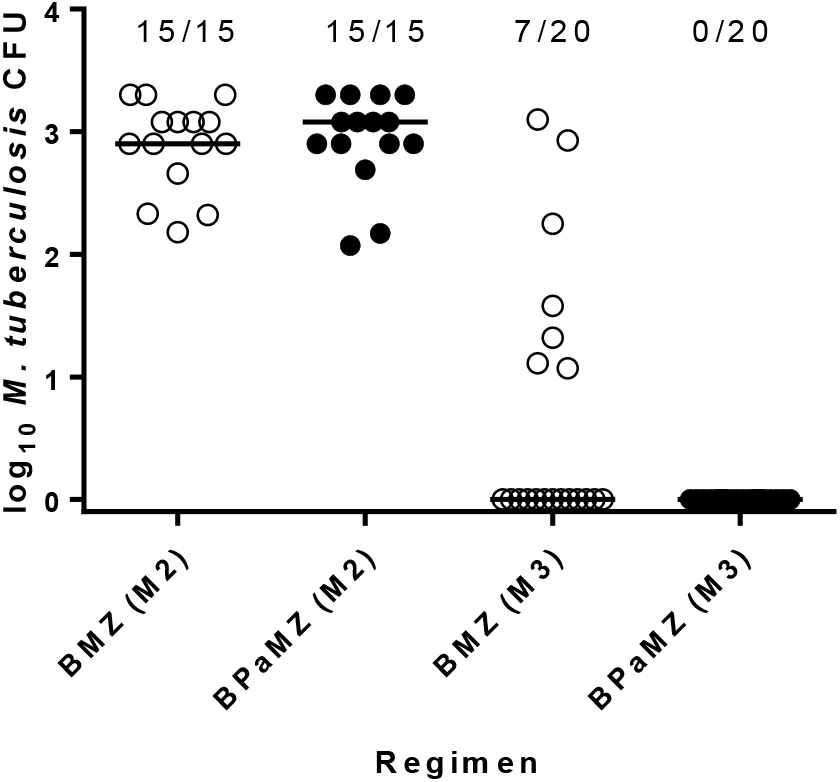
Proportion of relapses and individual mouse lung CFU counts (with median) after treatment of infection with *M. tuberculosis pncA* A146V mutant for two months (M2) and three months (M3) with each regimen. BMZ open black circles and BPaMZ solid black circles.

**Table 3.**
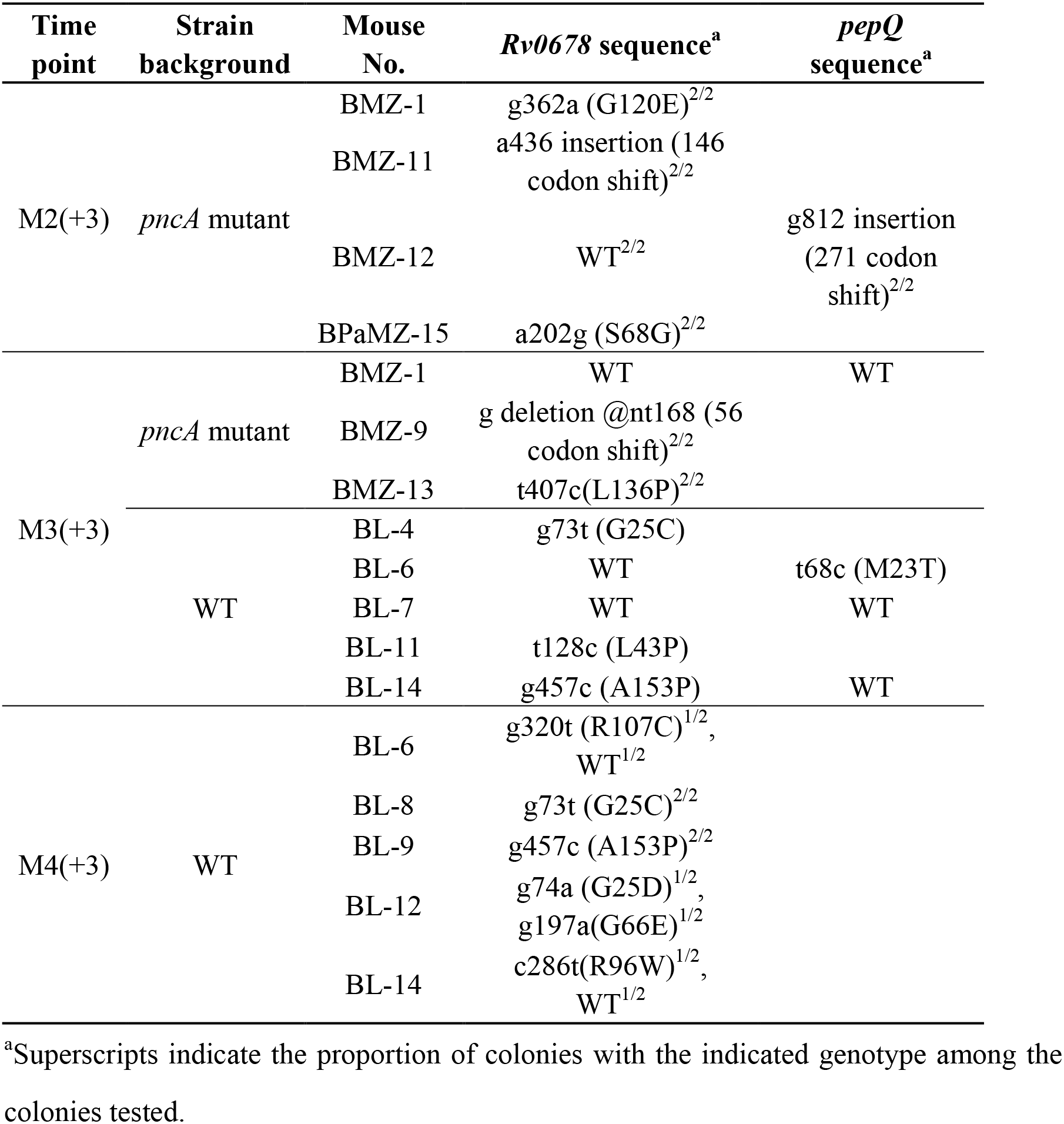
Mutations observed in *M. tuberculosis* colonies isolated from relapsing BALB/c mice on bedaquiline-containing plates in Experiment 2

Among mice infected with the H37Rv parent, addition of PMD to BL was associated with significantly lower mean CFU counts after 1 (*P* < 0.0001) and 2 (*P* = 0.0006) months of treatment (Table 2). Addition of PMD also had a marked effect on sterilizing activity. For example, the proportion of mice relapsing after treatment with BPaL for just 2 months (7/15 [47%)] was lower than that observed in mice receiving BL for 4 months (14/15 [93%]) (P = 0.01), indicating that inclusion of PMD reduced the treatment duration necessary to prevent relapse by at least 2 months. Addition of PMD also prevented the selection of BDQ-resistant mutants. Seven, five and five mice relapsing after receiving BL for 2, 3 and 4 months, respectively, had CFU growing on BDQ-containing plates exceeding 1% of the total CFU count and an additional mouse in the 2-month treatment cohort barely missed this threshold. The actual proportions of CFU growing on BDQ increased with treatment duration (1-5%, 2-22% and 2-41% after 2, 3 and 4 months of treatment, respectively). In contrast, no growth was observed on BDQ-containing plates in any mouse relapsing after BPaL treatment (*P* < 0.0001). Four of the 5 isolates from BDQ-containing plates at the M3+3 time point were tested and had mutations detected in *Rv0678* (3 isolates with g73t, t128c, or g457c substitutions) or *pepQ* (1 isolate with t68c substitution), while the remaining isolate had wild-type sequences in these genes (Table 3). All 5 mice relapsing at M4+3 harbored *Rv0678* mutants with single g320t, g73t, g457c, or c286t substitutions or both g74a and g197a substitutions in 2 isolates from one mouse (Table 3).

As expected, infection with the PMD-resistant *ddn* mutant eliminated the contribution of PMD to both the BPaMZ and BPaL regimens (Table 4). In fact, a trend towards modest dose-dependent antagonism was observed in mean CFU count comparisons when adding PMD to these combinations.

**Table 4.** Lung CFU counts assessed during treatment against a *ddn* mutant and proportion of mice relapsing after treatment completion in Experiment 2

| Regimen<sup>a</sup> | Mean lung log<sub>10</sub> CFU count (±SD)<sup>b</sup> |  |  |  | Proportion of mice relapsing after treatment for |  |  |
| --- | --- | --- | --- | --- | --- | --- | --- |
|  | D-14 | D0 | M1 | M2 | 1 mo. | 2 mo. | 3 mo. |
| Untreated | 4.23±0.07 | 7.61±0.19 |  |  |  |  |  |
| Pa<sub>50</sub> |  |  | 7.34±0.12 | <sup>c</sup> |  |  |  |
| Pa<sub>100</sub> |  |  | 7.26±0.07 | <sup>c</sup> |  |  |  |
| BL |  |  | 4.54±0.17 | 2.97±0.28 |  |  | 15/15 |
| BPa<sub>50</sub>L |  |  | 4.68±0.41 | 3.05±0.28 |  |  | 15/15 |
| BPa<sub>100</sub>L |  |  | 5.31±0.35 | 3.33±0.09 |  |  | 15/15 |
| BMZ |  |  | 2.36±0.63 | 0.00±0.00 | 15/15 | 2/15 |  |
| BPa<sub>50</sub>MZ |  |  | 2.49±0.24 | 0.08±0.19 | 15/15 | 1/15 |  |
| BPa<sub>100</sub>MZ |  |  | 2.73±0.46 | 0.00±0.00 | 15/15 | 1/15 |  |

<sup>a</sup>Drug regimen abbreviations are as follows: BL, bedaquiline + linezolid; BPaL, bedaquiline + pretomanid + linezolid; BMZ, bedaquiline + moxifloxacin + pyrazinamide; BPaMZ, bedaquiline + pretomanid + moxifloxacin + pyrazinamide.
<sup>b</sup>Time points are shown as days (D-14 or D0) or months (M1 or M2) of treatment. 1 mo. indicates that the mice were held for 3 additional months after completing 1 month of treatment.
<sup>c</sup>Five mice receiving Pa monotherapy at either 50 or 100 mg/kg became ill, were euthanized at Week 1, and had a mean lung CFU count of 9.07 log<sub>10</sub> CFU. One mouse in the BL group also died at Week 1, due to a gavage accident. The lung contained 7.04 log<sub>10</sub> CFU.

### Experiment 3. Comparison of BPaMZ and BMZ in C3HeB/FeJ mice

The scheme for this experiment is shown in Table S5. The two aerosol infections each implanted approximately 3 log_10_ CFU in the lungs of C3HeB/FeJ mice. By the start of treatment 4 weeks after the first infection, the mean CFU count had increased to 9.43±0.33 log_10_. Due to the unexpectedly high burden of infection and the rapidly evolving lung damage underway at treatment onset, substantial mortality was observed over the ensuing 2-month treatment period despite the strong bactericidal effect of both regimens. Addition of PMD to the BMZ regimen extended the median survival from 21 days to more than 60 days (P < 0.0001) (Figure 3) and significantly increased the bactericidal activity. After 1 month of treatment, the median lung CFU count was 2.4 log_10_ lower among mice receiving BPaMZ compared to BMZ (*P* < 0.01) (Figure 4). After 2 months of treatment, only 2 BMZ-treated mice survived compared to 10 BPaMZ-treated mice. Other than 1 BPaMZ-treated mouse with 2 CFU, all mice were culture-negative. No colonies were isolated on plates containing BDQ (0.06 μg/ml) or PMD (2 μg/ml) at either time point.

**Figure 3.**
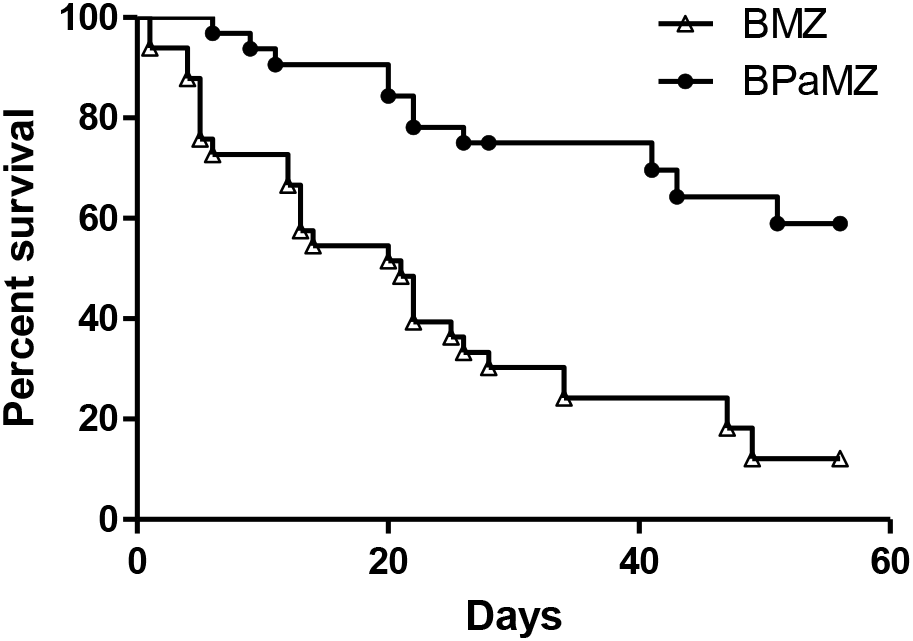
Survival of C3HeB/FeJ mice infected with *M. tuberculosis* HN878 from the onset of treatment with BMZ or BPaMZ.

**Figure 4.**
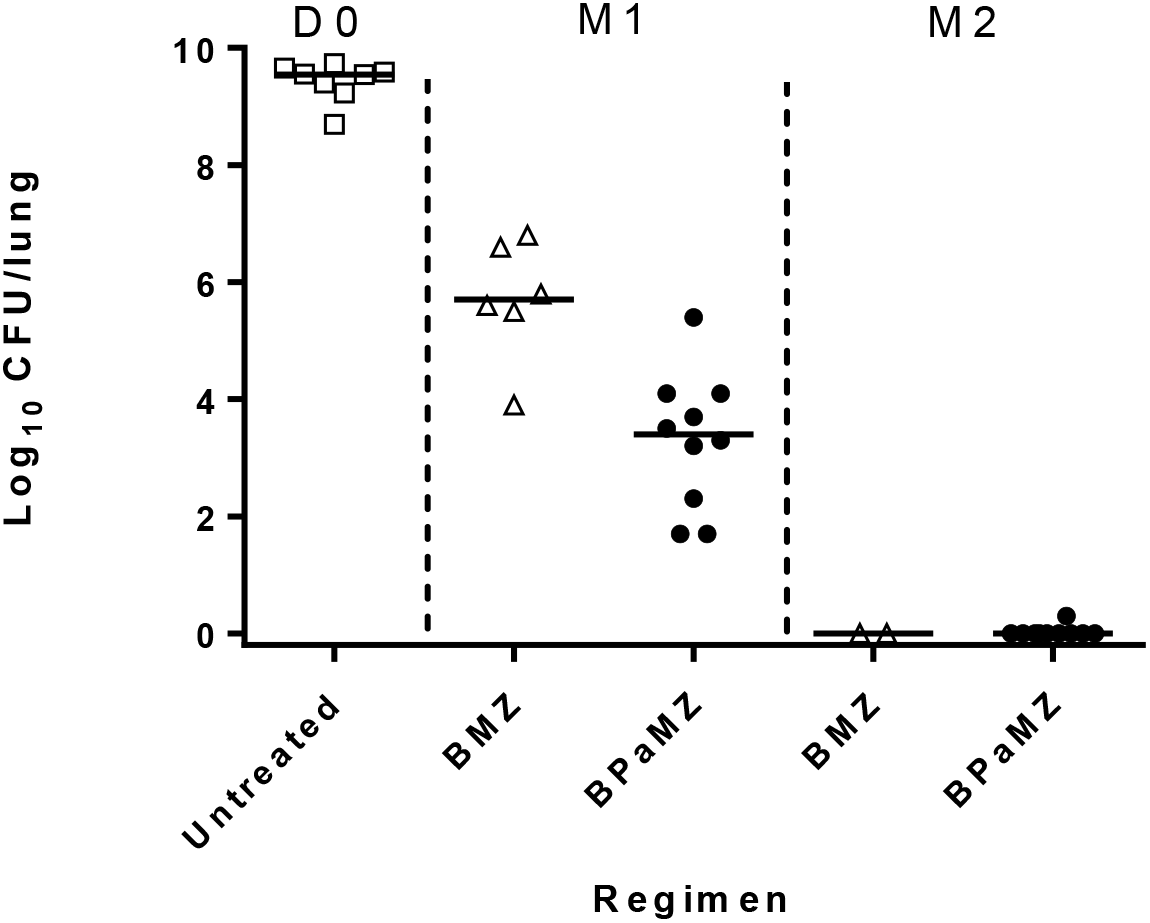
Lung CFU counts assessed during treatment in C3HeB/FeJ mice infected. Data points indicate individual mouse CFU counts. Horizontal black line indicates the median.

At 4 weeks post-infection (D0) with a high aerosol dose of *M. tuberculosis* HN878 under an accelerated disease protocol, C3HeB/FeJ mice exhibited extensive lung involvement with both cellular and caseating lesions (Figure 5). Cellular lesions were composed of neutrophilic clusters interspersed with lymphocytes and epithelioid macrophages. Caseating lesions included both isolated and coalescing granulomas with varying degrees of central caseation and cellularity (Figure 5, D0). Dense neutrophilic infiltration and abundant intracellular and extracellular acid-fast bacilli were evident at the foamy macrophage:caseum interface. By 6 weeks post-infection (W2 of treatment), the extent of lung disease had increased despite treatment. While some caseating lesions displayed more organized structures with an increasingly well-defined fibrous rim, more extensive central caseation and even cavitation, other areas displayed extensive infiltration with exudative pneumonitis (Figure 5 W2). Extracellular bacteria were increasingly evident in the acellular caseum. At 8 weeks post-infection (M1 of treatment), lung volumes were dominated by large areas of necrosis with central caseation and cavitary lesions (Figure 5 M1). Multiple lesion types presented at this time, suggesting heterogeneous disease progression. After 1 month of treatment with BPaMZ or BMZ, acid-fast staining was more diffuse and reflective of structural deterioration of bacteria by the highly bactericidal regimens. After 2 months of treatment, similar pathological changes were evident on H&E staining (Figure 5 M2), but no visible intact acid-fast bacilli were observed in either treatment group.

**Figure 5.**
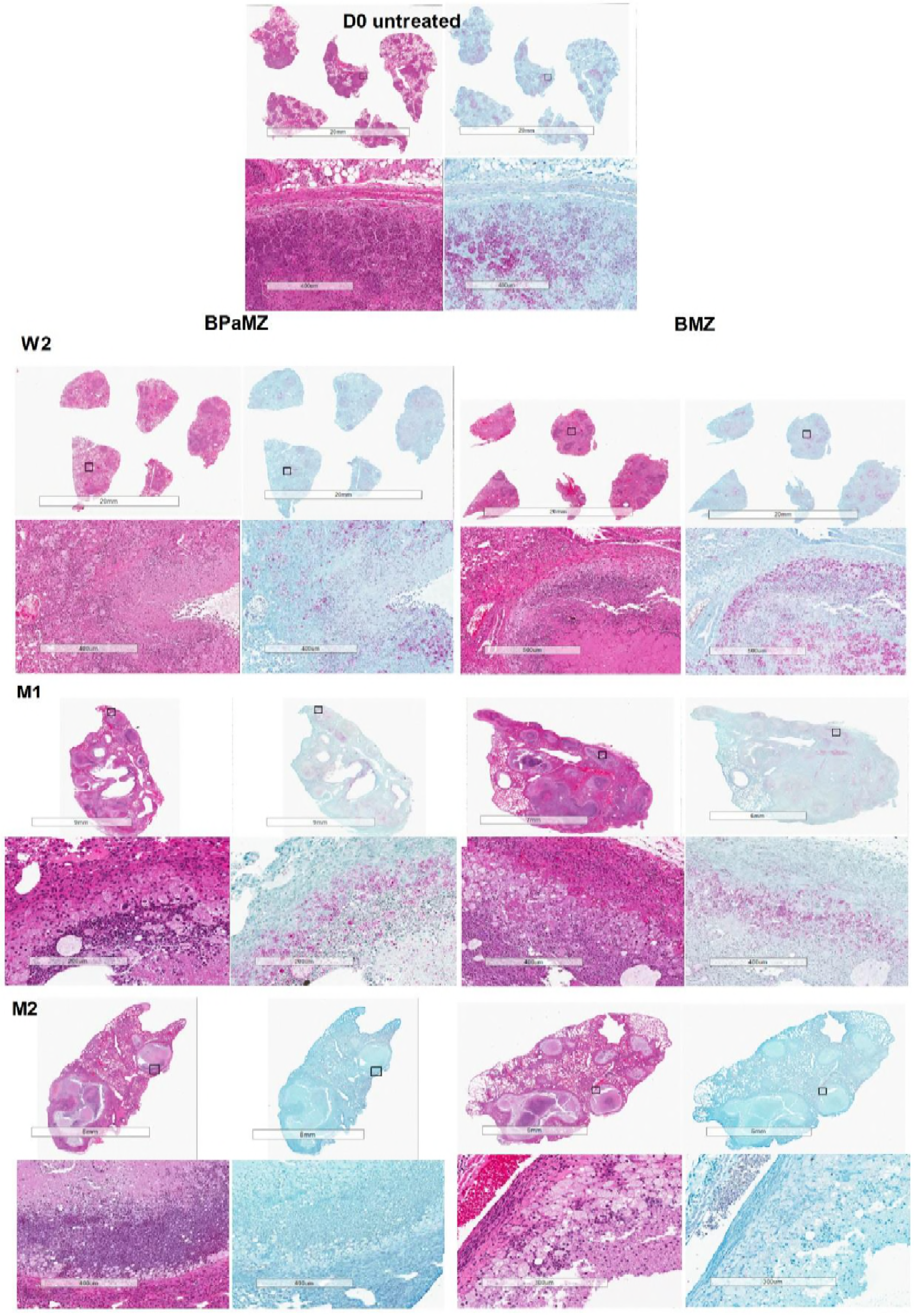
Lung histopathology in C3HeB/FeJ mice before and during treatment with BPaMZ (left) or BMZ (right) beginning 4 weeks post-infection with *M. tuberculosis* HN878. Hematoxylin and eosin (H&E) and Ziehl-Neelsen (AFB) staining was performed on lung tissue sections. Low-power views of an entire lung sectionsubmitted (upper) and higher-power views of individual granulomas or cavitary lesions (lower) from representative mice in each group are shown. D0, treatment initiation (4 weeks post-infection); W2, status post 2 weeks of treatment; M1, status post 1 month of treatment; M2, status post 2 months of treatment

## Discussion

The BPaL and BPaMZ regimens have the potential to transform the treatment of both drug-susceptible and drug-resistant TB. The former shows promise as an oral short-course (e.g., 6-month) regimen for multidrug- and extensively drug-resistant TB (7), while the latter may be capable of shortening the treatment of drug-susceptible TB to 4 months or less and could reasonably be expected to do the same for multidrug-resistant TB with preserved susceptibility to MXF and PZA(5). Both regimens were identified in a comprehensive screening program seeking to identify broad-spectrum regimens containing two or more novel agents with minimal pre-existing resistance in our high-dose aerosol infection model in BALB/c mice (4, 6). However, while the independent contributions of each other component of these regimens has been confirmed in this model (4, 6), the contribution of the investigational new drug PMD to the BPaMZ regimen was not previously demonstrated and its contribution to the BPaL regimen was only assessed as far as the bactericidal activity of the regimen over the first 2 months of treatment and not PMD’s contribution to the regimen’s sterilizing activity.

The present study assessed the contribution of PMD to the BPaMZ regimen in 3 different murine models of TB and its contribution to the sterilizing activity of BPaL in BALB/c mice. Studies using the relapse endpoint in BALB/c mice have demonstrated utility for regimen selection and estimation of treatment duration (25, 26). The present results reinforce our prior findings that BPaMZ has remarkable bactericidal and sterilizing activity in BALB/c mice (4) and extend them by demonstrating the independent contribution of PMD, as demonstrated by the BPaMZ regimen’s superior reduction of M1 lung CFU counts compared to BMZ in Experiment 1 and it superior prevention of relapse in Experiment 2. The present study also confirmed the contribution of PMD to the bactericidal activity of the BPaL regimen observed in prior studies (6, 12) and shows, for the first time, PMD’s key contribution to the sterilizing activity of this regimen.

Prior work has established immunocompromised athymic nude mice as a more stringent model for measuring the sterilizing activity of rifamycin-based regimens, as well as such regimens’ ability to restrict the emergence of resistance, indicating a beneficial contribution of cell-mediated immunity in the treatment response (15, 16). Therefore, it was expected that the eradication of cultivable bacteria with BPaMZ would require a longer duration of treatment in this strain. Nevertheless, BPaMZ rendered nude mice culture-negative with ≤ 2 months of treatment and prevented relapse in nearly 95% of mice after 2.5 months of treatment. The absence of relapse in nude mice is likely to reflect true sterilization of infection by the regimen, especially considering the rapid demise of 3 of 4 relapsing mice in the BMZ arm, and affirms the intrinsic sterilizing activity of this drug combination. It should be noted that the bactericidal and sterilizing activity of BPaMZ in this experiment was superior to that of a rifapentine-isoniazid-PZA regimen previously evaluated in the same model (15). Inclusion of PMD significantly reduced lung CFU counts over the first month of treatment and reduced the proportion of mice relapsing, although the effect of PMD on relapse did not reach statistical significance. These results suggest that PMD’s contribution to the regimen will extend to immunocompromised hosts.

To further examine the contribution of PMD to BPaMZ, we used the emerging C3HeB/FeJ mouse model to better mimic the pathophysiological conditions found within caseating human lung lesions (e.g., hypoxia, more neutral pH) (18–20, 22, 24, 27). Prior observations have indicated reduced diffusion of BDQ into the caseous regions of necrotic lung lesions relative to the bordering cellular regions and reduced activity of PZA in caseum with near-neutral pH in this strain (23, 24). On the other hand, PMD appears to diffuse well through caseum and is active under hypoxic conditions (10, 28), so should lend important bactericidal activity in caseous lesions. The high infectious dose and repeated aerosol infection protocol used in Experiment 3 resulted in extensive lung involvement with large, often coalescing, caseating granulomas, caseous pneumonia and, over time, cavitation. The massive bacterial burden and marked severity of lung disease present at the onset of treatment resulted in further pathologic progression and death despite the initiation of the highly bactericidal BPaMZ and BMZ regimens. Despite this “worst case scenario”, the inclusion of PMD in the regimen significantly increased the median survival time and the bactericidal activity, and the regimen rendered all mice culture-negative after 2 months of treatment, with the exception of a single mouse with 1 detectable CFU. It is tempting to speculate that the larger effect of PMD on M1 CFU counts in this model compared to BALB/c and nude mice was a result of its superior distribution and/or activity within caseating lesions compared to BDQ and PZA. Further studies are warranted to assess PMD’s independent treatment-shortening effect in this model. Nevertheless, these studies provide further reassurance that the results observed with BPaMZ, and the contribution of PMD specifically, are translatable to human TB.

BDQ is a key bactericidal and sterilizing component of the BPaMZ and BPaL regimens (4, 6). As such, it exerts strong selection pressure and bactericidal and sterilizing companion drugs are necessary to restrict the selective amplification of spontaneous BDQ-resistant mutants. Previous studies have identified BDQ-resistant mutants selected *in vitro*, in mice and in TB patients (13, 29–31). In most cases in which it emerged *in vivo*, resistance was attributable to mutations in the transcriptional repressor *Rv0678* or in the predicted proline aminopeptidase *pepQ*, although the latter mutation target has yet to be confirmed in BDQ-resistant clinical isolates. In the present study, we tracked the selection of BDQ-resistant mutants, and the ability of PMD to prevent such selection, in immunocompromised nude mice in Experiment 1, BALB/c mice in Experiment 2, and severely diseased C3HeB/FeJ mice in Experiment 3. Fortunately, the BPaMZ regimen proved to be quite robust to the emergence of resistance in each model. The only instance in which a BDQ-resistant isolate was observed after BPaMZ treatment was a single mouse in Experiment 2 that was originally infected with a PZA-resistant strain, compared to 6 mice receiving BMZ against the same strain. Likewise, inclusion of PMD in the BPaL regimen significantly reduced the proportion of relapsing mice with BDQ resistance. These findings demonstrate the limited ability of LZD and MXF, respectively, to prevent the selective amplification of BDQ resistance on their own and suggest that PZA may play an important role in preventing such amplification among phenotypic sub-populations similar to those represented in BALB/c mice.

Interestingly, no selection of BDQ resistance was observed with BPaMZ or BMZ treatment of C3HeB/FeJ mice despite the large bacterial burden and the expected limited contribution of PZA in caseous lesions in this model (24). Here, PMD and MXF may be especially effective in the caseum compartment while PZA again works in the intracellular compartment. Similarly, little BDQ resistance selection was observed in nude mice, and only in a BMZ-treated mouse, despite evidence of spontaneous resistant mutants present at baseline, when all nude mice sampled harbored *Rv0678* mutant sub-populations and 2 of 5 harbored spontaneous *pepQ* mutants (40%). Consistent with some prior observations (30, 32), frameshift mutations in *Rv0678* caused higher MICs than single nucleotide polymorphisms in *Rv0678* and *pepQ* mutations. It is noteworthy that the MICs associated with the latter mutants, in particular, hover around the recently proposed critical concentration for BDQ susceptibility testing on 7H11 agar (33) and hence may not be recognized as resistant despite their apparent reduced BDQ susceptibility and propensity for selective amplification despite combination therapy in mice.

In conclusion, PMD contributed significantly to the efficacy of both the BPaMZ and BPaL regimens and reduced the selection of bedaquiline-resistant mutants. These results support further clinical trials to confirm the therapeutic utility of each of these PMD-containing regimens.

## Methods

### Bacterial strains

*M. tuberculosis* H37Rv was mouse-passaged, frozen in aliquots, and subcultured in Middlebrook 7H9 broth supplemented with 10% oleic acid-albumin-dextrose-catalase (OADC) (Fisher, Pittsburgh, PA) and 0.05% Tween 80 prior to infection. An isogenic PZA-resistant mutant was selected from a mouse infected with the H37Rv strain and treated with PZA monotherapy and an isolated mutation in *pncA* (A146V) was confirmed by whole genome sequencing (24). The PZA MIC for this *pncA* mutant is ≥ 900 μg/ml (vs. 150 μg/ml for the parent H37Rv strain) by 7H9 broth dilution at pH 6.8. An isogenic PMD-resistant mutant was selected from a mouse infected with the H37Rv strain and treated with PMD monotherapy and an isolated mutation in *ddn* (M1T) was confirmed by whole genome sequencing. The pretomanid MIC for this *ddn* mutant is ≥ 16 μg/ml (vs. 0.06-0.125 μg/ml for the parent H37Rv strain) by 7H9 broth dilution. *M. tuberculosis* HN878 was used as frozen stocks prepared from a log-phase culture in Middlebrook 7H9 broth after mouse passage and was diluted in phosphate buffered saline (PBS) before infection.

### Aerosol infection with *M. tuberculosis*

All animal procedures were approved by the Animal Care and Use Committee of Johns Hopkins University. Six-week-old female BALB/c mice and immunodeficient athymic CD-1 nude mice (Charles River Laboratories, Wilmington, MA) were infected with *M. tuberculosis* H37Rv or the isogenic *pncA* or *ddn* mutant, using the Inhalation Exposure System (Glas-Col, Terre Haute, IN) and a fresh log-phase broth culture (optical density at 600 nm, 0.8 to 1.0), with the goal of implanting 4 log_10_ CFU in the lungs of each mouse (4). Four or five mice were humanely killed 1 day after infection (D-13) and on the day of treatment initiation (D0) to determine the number of bacteria implanted in the lungs and at the start of treatment, respectively.

Female C3HeB/FeJ mice (Jackson Labs, Bar Harbor, ME), 10 weeks old, were aerosol-infected with *M. tuberculosis* HN878 on two occasions spaced 10 days apart (with mice divided into 2 runs per occasion) in a repeated infection protocol intended to promote more advanced caseating lung lesions. On each occasion, a frozen stock culture was thawed and diluted with the intention to implant approximately 200 CFU per run. Treatment started at 4 weeks after the first infection. Six and nine mice (2 or 3 mice per run) were sacrificed for lung CFU counts on W-4 and D0 to determine the number of CFU implanted and the number present at the start of treatment, respectively.

### Chemotherapy

Drugs were prepared as previously described and administered once daily, 5 days per week, by gavage (11). The drug doses (in mg/kg) were as follows: BDQ, 25; PMD, 100; MXF, 100; PZA, 150; LZD, 100 (4, 6). BDQ and PMD were prepared separately and administered together after mixing just prior to administration. MXF, MXF+PZA or LZD were administered at least 4 hours later.

### Assessment of treatment efficacy

Efficacy was assessed on the basis of lung CFU counts at selected time points during treatment (a measure of bactericidal activity) and the proportion of mice with culture-positive relapse after treatment completion (a measure of sterilizing activity). Lung homogenates were plated in serial 10-fold dilutions onto selective 7H11 agar plates supplemented with 0.4% activated charcoal to reduce carryover effects (11) and incubated for 6 weeks before determining final CFU counts.

### Evaluation of resistance selection

Aliquots representing one-fifth of the lung homogenates were plated directly onto selective 7H11 agar containing 0.06 or 0.125 μg/ml of BDQ or 2 μg/ml of PMD to quantify the proportion of CFU resistant to either drug at selected time points before, during and after treatment. Colonies isolated on BDQ-containing plates were selected and analyzed by PCR and DNA sequencing of the *rv0678, pepQ*, and *atpE* genes, as described previously (13). The MICs of bedaquiline-resistant isolates and H37Rv were determined using the broth macrodilution method with doubling concentrations of bedaquiline from 0.03 to 2 μg/ml (13). Briefly, tubes containing 2.5 ml of 7H9 broth plus OADC without Tween 80 with the above-mentioned concentrations of bedaquiline were inoculated with 10^5^ CFU of log-phase culture of H37Rv or BDQ-resistant isolates. The MIC was defined as the lowest concentration that prevented visible growth after 14 days of incubation at 37°C.

### 16S ribosomal RNA sequencing for identification of bacteria

Genomic DNA was extracted from bacterial colonies using the cetyltrimethylammonium bromide (CTAB) method. 16S rRNA was amplified with primers 16S-F (5’-AGAGTTTGATCCTGGCTCAG-3’) and 16S-R (5’-ACGGGCGGTGTCTACAA-3’) targeting positions 11–1399 of the 16S rRNA gene (34). PCR products were purified with QIAquick PCR purification kits (QIAGEN, Germany), mixed with primers, then sequenced (GeneWiz Inc.). Species identification was performed using BLAST search (GenBank database sequences) with sequence data.

### Statistical analysis

Lung CFU counts (x) were log-transformed (as x+1) before analysis, and mean and median CFU counts were compared using Student’s t test and Kruskal-Wallis tests, respectively. The proportions of mice relapsing were compared using Fisher’s Exact test. Survival analyses were performed using the Kaplan-Meier method (20), and the log rank test was used to compare the observed differences in survival. All analyses were performed with GraphPad Prism version 5 (GraphPad, San Diego, CA).

## Acknowledgements

This research was supported by TB Alliance with support from Australia Aid, the Bill and Melinda Gates Foundation, the Germany Federal Ministry of Education and Research through KfW, Global Health Innovative Technology Fund, Irish Aid, Netherlands Ministry of Foreign Affairs, UK Aid and the UK Department of Health; and the National Institutes of Health (R01-AI111992 to E.N.).

